# A high-throughput, automated, cell-free expression and screening platform for antibody discovery

**DOI:** 10.1101/2021.11.04.467378

**Authors:** Andrew C. Hunt, Bastian Vögeli, Weston K. Kightlinger, Danielle J. Yoesep, Antje Krüger, Michael C. Jewett

## Abstract

Antibody discovery is bottlenecked by the individual expression and evaluation of antigen-specific hits. Here, we address this gap by developing an automated workflow combining cell-free DNA template generation, protein synthesis, and high-throughput binding measurements of antibody fragments in a process that takes hours rather than weeks. We apply this workflow to 119 published SARS-CoV-2 neutralizing antibodies and demonstrate rapid identification of the most potent antibody candidates.

## Main Body

Antibodies are widely used as protein-based drugs and diagnostics. They are the critical component in immunoassays enabling rapid diagnostics^1^ and constitute one of the fastest-growing classes of therapeutics with nearly 25% of new FDA-approved drugs in 2020 being antibodies^2,3^. Antibodies have also recently garnered attention as potential countermeasures for emerging pathogens, and currently constitute the majority of emergency use authorized treatments for COVID-19 that inhibit the SARS-CoV-2 virus^4–6^.

Modern workflows for antibody discovery utilize either directed evolution or the isolation of single B-cell clones from convalescent patients or animals to go from >10^8^ possible sequences to a pool of ~10^3^ candidates targeting the desired antigen. However, once this pool of candidates has been generated, state-of-the-art workflows still rely on labor-intensive and poorly scalable procedures (e.g., plasmid-based cloning, transfection, cell-based protein expression, protein purification, binding assessment through enzyme-linked immunosorbent assays (ELISAs), etc.) to individually evaluate and identify the best antibody candidates^7,8^. These labor-intensive procedures take weeks to months and represent a major bottleneck in antibody discovery. The effort to identify antibodies against emerging threats like the SARS-CoV-2 during the COVID-19 pandemic has highlighted (i) the importance of rapid and high-throughput antibody discovery platforms and (ii) the importance of identifying high-affinity antibodies targeting conserved epitopes^9,10^ or non-overlapping epitopes^11,12^ to resist viral escape and increase the ability to neutralize viral variants^13,14^; both of which have required intensive screening campaigns. A further challenge is that existing antibody discovery processes frequently have low efficiency, with very few of the screened candidates being potent neutralizers in the case of SARS-CoV-2 (Supplementary Table 1). Taken together, these limitations in existing antibody discovery processes suggest the urgent need for faster and higher throughput screens.

Cell-free protein synthesis (CFPS)^15,16^, the manufacture of proteins without living cells using crude extracts or purified components, is an attractive tool to overcome these limitations. Towards this goal, a variety of CFPS systems for antibody expression have been developed^17–23^. However, to our knowledge, an end-to-end (DNA to data) automatable antibody screening workflow combining CFPS with a high-throughput protein-protein interaction screen has yet to be developed.

Here we describe such an integrated pipeline for antibody expression and evaluation to address critical screening limitations in current antibody discovery pipelines. The workflow leverages four key developments (Fig. 1a): (i) DNA assembly and amplification methods that do not require living cells, (ii) CFPS systems that can work directly from linear DNA templates and can generate disulfide-bonded antibody molecules, (iii) an Amplified Luminescent Proximity Homogeneous Linked Immunosorbent Assay (AlphaLISA) that enables rapid protein-protein interaction (PPI) or binding characterization without protein purification^24^, and (iv) robotic and acoustic liquid handling that enables a highly parallel and miniaturized workflow. Our integrated workflow is end-to-end automatable and enables a single researcher to express and profile the antigen-specific binding of hundreds of antibodies in 24 hours. As a model, we applied our workflow to profile a diverse set of 120 previously published antibodies, 119 of which are antibodies targeting the SARS-CoV-2 spike glycoprotein (S trimer). These antibodies were selected based on the availability of sequence, structural, SARS-CoV-2 neutralization, and binding information, with 84 being drawn from Brouwer *et al.*^25^ and the remainder from diverse sources^9,26–40^ (Supplementary Table 2 and 3). The antibodies span four orders of magnitude in neutralization potency and target a variety of domains and epitopes.

**Fig. 1.**
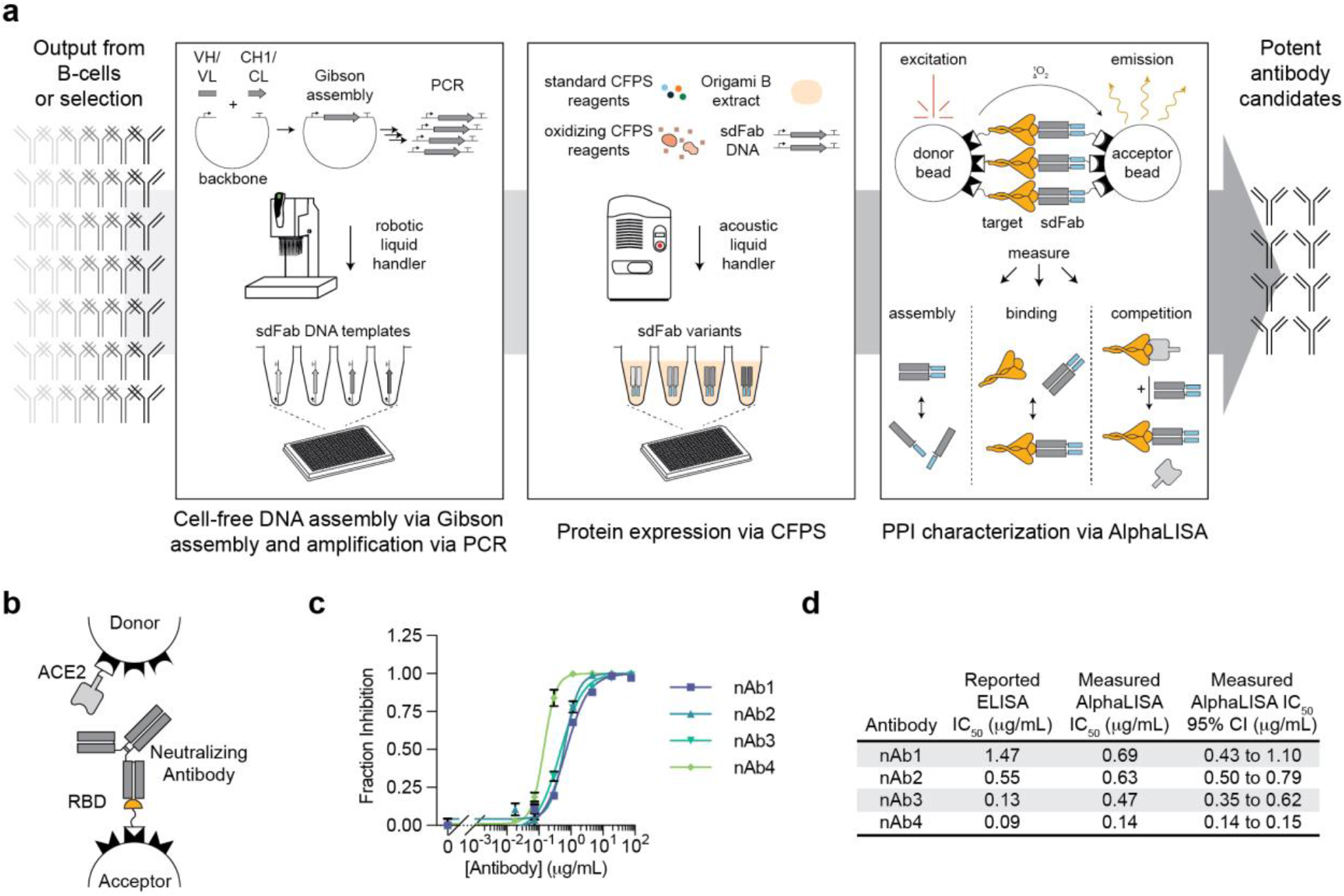
A high-throughput, cell-free antibody screening workflow. **a,** Schematic of the steps involved in the cell-free antibody screening workflow. **b**, Diagram of the AlphaLISA screen for neutralizing antibodies via competition with ACE2 for the SARS-CoV-2 RBD. **c**, Evaluation of commercial neutralizing antibodies (nAbs) in the AlphaLISA ACE2 competition screen (n=3 independent replicates ± SEM). **d**, Comparison of the reported and measured potencies of commercial neutralizing antibodies.

We first implemented a cell-free method for DNA assembly and amplification by adapting and optimizing recently reported protocols for high-throughput construction of DNA templates for CFPS^17,19,41,42^. The method consists of a Gibson assembly step, followed by PCR amplification of the linear expression template (LET) using the unpurified Gibson assembly product as a template. The key idea was to create a versatile approach for rapid construction of DNA templates without the requirement of cell culture, allowing DNA assembly and amplification in less than 3 hours entirely in 384 well plates. To validate the method, we applied it to the assembly and amplification of a LET for sfGFP expression and only observed sfGFP expression in the presence of properly assembled DNA template (Supplementary Fig. 1a-c). To assemble antibody DNA templates, we purchased synthetic, double-stranded linear DNA coding for the desired variable heavy (VH) and variable light (VL) chain sequences. These DNAs were assembled with DNA coding for the appropriate heavy chain constant (CH1) or light chain constant (CL) antigen-binding fragment (Fab) domains in addition to a separate piece of DNA coding for the backbone of the pJL1 vector. These sequences were subsequently amplified by PCR to generate LETs (Supplementary Fig. 1d-f). Previous works suggest that this workflow could be compatible with PCR products amplified from single B-cells from an immunized animal^17,19,41^. In addition to being fast, this workflow also affords flexibility, allowing assembly of different antibody formats (e.g., full-length, Fab, sdFab) containing different purification or immobilization tags by using different antibody constant regions in the assembly reaction.

We next demonstrated rapid antibody expression in a crude *E. coli* based CFPS system. We developed a high-yielding (1,390 ± 32 μg/mL sfGFP, Supplementary Fig. 1c) crude *E. coli* lysate-based CFPS system from the Origami™ B(DE3) strain (Supplementary Fig. 2), which contains mutations in the *E. coli* reductase genes *trxB* and *gor* to enable the formation of disulfide bonds in the cytoplasm^43^. By pretreating the extract with the reductase-inhibitor iodoacetamide (IAM) to further stabilize the redox environment^44–46^ and supplementing the reaction with purified *E. coli* disulfide bond isomerase DsbC and prolyl isomerase FkpA^20,47,48^, we successfully expressed and assembled full-length trastuzumab, a model anti-HER2 antibody^49^, from linear DNA templates (Supplementary Fig. 2a). However, like others^17–21^, we found that the efficient assembly of full-length antibodies in CFPS can require further optimization (e.g., temperature, DNA template ratio, DNA template expression timing) which is not optimal for high-throughput screening. Like reports by Ojima-Kato *et al.*^17–19^, we found that the assembly of synthetically dimerized antigen-binding fragments (sdFab, also called ecobodies^17,19^ or zipbodies^18^) were more consistent than their corresponding standard Fabs in CFPS for a small panel of antibodies and opted to utilize the sdFab format for expression (Supplementary Fig 2b-c). Using acoustic liquid handling we can assemble CFPS reactions to express each sdFab variant from cell-free assembled and amplified DNA in 384-well plates (Fig. 1a).

Following DNA assembly and CFPS, antigen-specific binding was evaluated. To characterize the PPIs of the expressed sdFab antibody candidates, we developed an AlphaLISA method to characterize PPIs directly from CFPS reactions. AlphaLISA is an in-solution and wash-free assay that is designed for high-throughput screening and is compatible with crude cell-lysates^24^. In AlphaLISA, non-covalent capture chemistries are used to immobilize the proteins of interest on donor and acceptor beads, which generate a chemiluminescent signal when in proximity of one another and excited by a 680 nm laser. We developed AlphaLISA methods to enable the measurement of both direct binding to an antigen as well as competition for specific epitopes. We first sought to validate that AlphaLISA is tolerant of crude CFPS reactions. We observed that CFPS does not interfere with the measurement chemistry (Supplementary Fig. 3a), but that certain reaction components can disrupt protein immobilization to the bead which can be circumvented with the appropriate choice of immobilization chemistry (Supplementary Fig 3b-c). We found that the Ni-Chelate beads were not tolerant of the high salt concentrations and high concentration of histidine present in CFPS, likely due to charge screening and Ni chelation respectively hindering immobilization of the hisx6 tagged protein. To validate the ability of AlphaLISA to profile neutralizing antibodies, we tested the ability of five different commercial antibodies to compete with the SARS-CoV-2 target human receptor Angiotensin-Converting Enzyme 2 (ACE2) for binding of the SARS-CoV-2 Receptor Binding Domain (RBD) and found that our determined rank order of IC_50_ values aligns with the reported ELISA IC_50_’s (Fig. 1b–d). Further, we utilized AlphaLISA to develop a sdFab assembly screen to monitor antibody expression and assembly in CFPS, a laborious step that traditionally requires SDS-PAGE. The measurement immobilizes the heavy and light chains of the sdFab to the AlphaLISA beads, resulting in signal when the two chains are assembled (Supplementary Fig. 3d). The AlphaLISA assembly assay shows consistent prediction of antibody assembly with SDS-PAGE on a panel of sdFabs and can thus be used to identify when sdFab expression or assembly fails (Supplementary Fig. 3e).

Using the developed workflow, we next evaluated a set of 120 unique antibodies using AlphaLISA to measure antibody binding to the SARS-CoV-2 S trimer, binding to the SARS-CoV-2 RBD, competition with ACE2 for RBD binding, and assembly of their heavy and light chains in CFPS (Fig. 1a and Fig. 2). Antibodies were expressed and evaluated in triplicate. AlphaLISA replicates were found to be consistent with one another, validating that the acoustic liquid handling workflow is robust (Supplementary Fig. 4). Samples were evaluated for significant assembly, binding to, or competition with a given target using a two-sided student’s t-test corrected for multiple testing using the Benjamini and Hochberg False Discovery Rate procedure (FDR)^50^. Within the diverse set of 36 antibodies, we observed assembly for 36 out of 36 tested antibodies, S trimer binding for 28 out of 35 antibodies reported to bind the S trimer, RBD binding for 23 out of the 34 antibodies reported to bind the RBD, and ACE2 competition for 16 out of 31 antibodies reported to compete with ACE2 (Fig. 2a, Supplementary Fig. 5). For the set of 84 antibodies from Brouwer *et al.,* we observed assembly of 80 out 84 antibodies and binding to the S trimer and RBD for many of the antibodies that showed strong binding via ELISA (Fig. 2b–d). We compared ACE2 competition against neutralization since it has been reported that more than 90% of neutralizing antibodies block the RBD and ACE2 interaction^28,51^ and similar competition assays have been reported to correlate with neutralization potency^28,52^ (Fig. 2e). We observed ACE2 competition, as well as strong S trimer and RBD binding, for 4 out of 5 antibodies reported to compete with ACE2, which also represent the four most potent neutralizers in the Brouwer *et al.* data set.

**Fig. 2.**
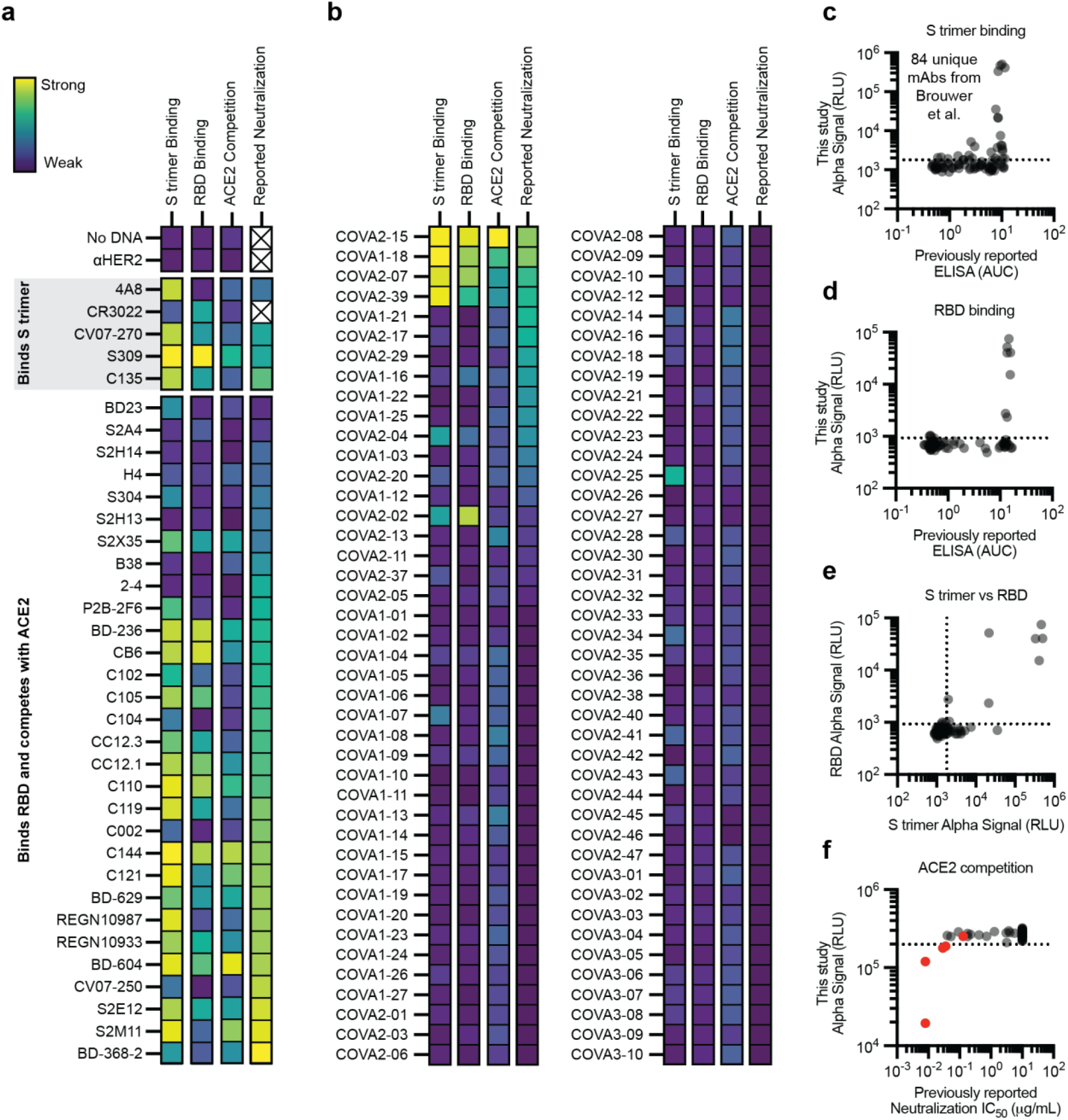
Performance of the cell-free antibody screening workflow evaluated on SARS-CoV-2 neutralizing antibodies. **a-f**, AlphaLISA data are presented as the mean of 3 independent replicates. A dashed line indicates three standard deviations away from the background signal. **a-b**, Heatmap of the binding of previously published antibodies measured using AlphaLISA to detect S trimer binding (log_10_ scaled), RBD binding (log10 scaled), and ACE2 competition (linearly scaled). AlphaLISA data are presented as the mean of 3 independent replicates. The lowest reported neutralization IC_50_ value is also plotted for comparison (log10 scaled) and an X indicates no relevant data available (Supplementary Table 2). **a** Heatmap of the binding of 36 diverse antibodies. **b**, Heatmap of the binding of all 84 antibodies in the Brouwer *et al.* data set. **c-d**, Parity plots comparing the AlphaLISA the 84 antibodies in the Brouwer *et al*. data set vs the published ELISA data. A dashed line indicates three standard deviations away from the background. **c**, S trimer binding. **d**, RBD binding. **e**, Comparison of the S trimer and RBD AlphaLISA binding data. **f**, Parity plot comparing the AlphaLISA ACE2 competition data for the 84 antibodies in the Brouwer *et al.* data set vs the published pseudovirus neutralization data. Antibodies that were reported to compete with ACE2 by Brouwer *et al.* are plotted in red.

Notably, we observed ACE2 competition for 10 out of 13 antibodies in the overall data set whose neutralization IC_50_ values are less than 0.01 μg/mL. While some less-effective neutralizers could not be completely characterized in our screen, we consistently identified potent neutralizing antibodies in our rapid cell-free screening workflow whose mechanism is ACE2 competition. Consistent with their binding specificities, we observed that 4A8, an n-terminal domain targeted antibody^37^, only showed strong interaction with the S trimer and that CR3022, whose target epitope is occluded in the S trimer^34,53^, showed binding to the RBD, but weak binding to the S trimer. Surprisingly, the S309 antibody in the sdFab format exhibited competition with ACE2 although it has been previously reported not to compete with ACE2^9^, which will require further study. Taken together, the binding and competitive AlphaLISA data generated by our workflow are self-consistent and largely align with the literature (Supplementary Table 4). Further improvements to the dynamic range of the PPI measurements could broaden the utility for performing antigenic mapping of the immune response to antigens. Inclusion of other binding targets could allow researchers to easily evaluate targeting to different domains (e.g., SARS-CoV-2 N-terminal domain) or look for antibodies targeting conserved epitopes by evaluating cross-reactivity with other related viruses (e.g., SARS-CoV, etc.).

In summary, we developed an integrated and automated workflow for antibody screening by combining methods for cell-free DNA assembly and amplification, cell-free protein expression, and highly parallel binding characterization via AlphaLISA. This workflow has two key features. First, it is fast. The entire workflow for all 120 antibodies evaluated in this study was completed in triplicate in less than 24 hours in two consecutive working days by a single researcher, highlighting the workflow’s speed and throughput. Second, integration of the AlphaLISA assay in cell-free extracts without the need for protein purification facilitates direct evaluation of synthesized antibodies in high-throughput. This is important because this is frequently the limiting step in previously published methods. Looking forward, we anticipate that the increased speed and throughput afforded by our workflow will enable researchers to easily and rapidly screen thousands of antibodies, facilitating down-selection to a few highly potent candidates that can be expressed at larger scales in cells or using CFPS and subjected to deeper developability testing. In this way, our method is poised to aid in the discovery of medical countermeasures in future pandemics, and more broadly, in the development of antibodies for therapeutic, diagnostic, and research applications.

## Acknowledgments

We thank Ashty S. Karim (ORCID 0000-0002-5789-7715) for discussions about research direction and comments on the manuscript and Kosuke Seki for the donation of the fLuc plasmid. M.C.J. acknowledges support from the David and Lucile Packard Foundation, the Camille Dreyfus Teacher-Scholar Program, and the Defense Threat Reduction Agency Grants HDTRA1-15-10052, HDTRA-12-01-0004, and HDTRA-12-11-0038. A.C.H. was supported by a Department of Defense National Defense Science and Engineering Graduate (NDSEG) Fellowship (NDSEG-36373).

## Author Contributions

A.C.H., B.V., and M.C.J. conceived of the study. A.C.H., B.V., and A.K.G. completed the research. W.K.K. and D.J.Y. assisted with cell-free expression methods. M.C.J. supervised the research. A.C.H. wrote the manuscript. All authors commented on and edited the manuscript.

