## Supplemental Materials for "A high-throughput, automated, cell-free expression and screening platform for antibody discovery"

**This PDF file includes:**

Supplementary Figures 1 to 5
Supplementary Table 1
Captions for Supplementary Tables 2 to 5
Materials and Methods

**Other Supplementary Materials for this manuscript include the following:**

Supplementary Tables 2 to 5 as separate .xlsx files

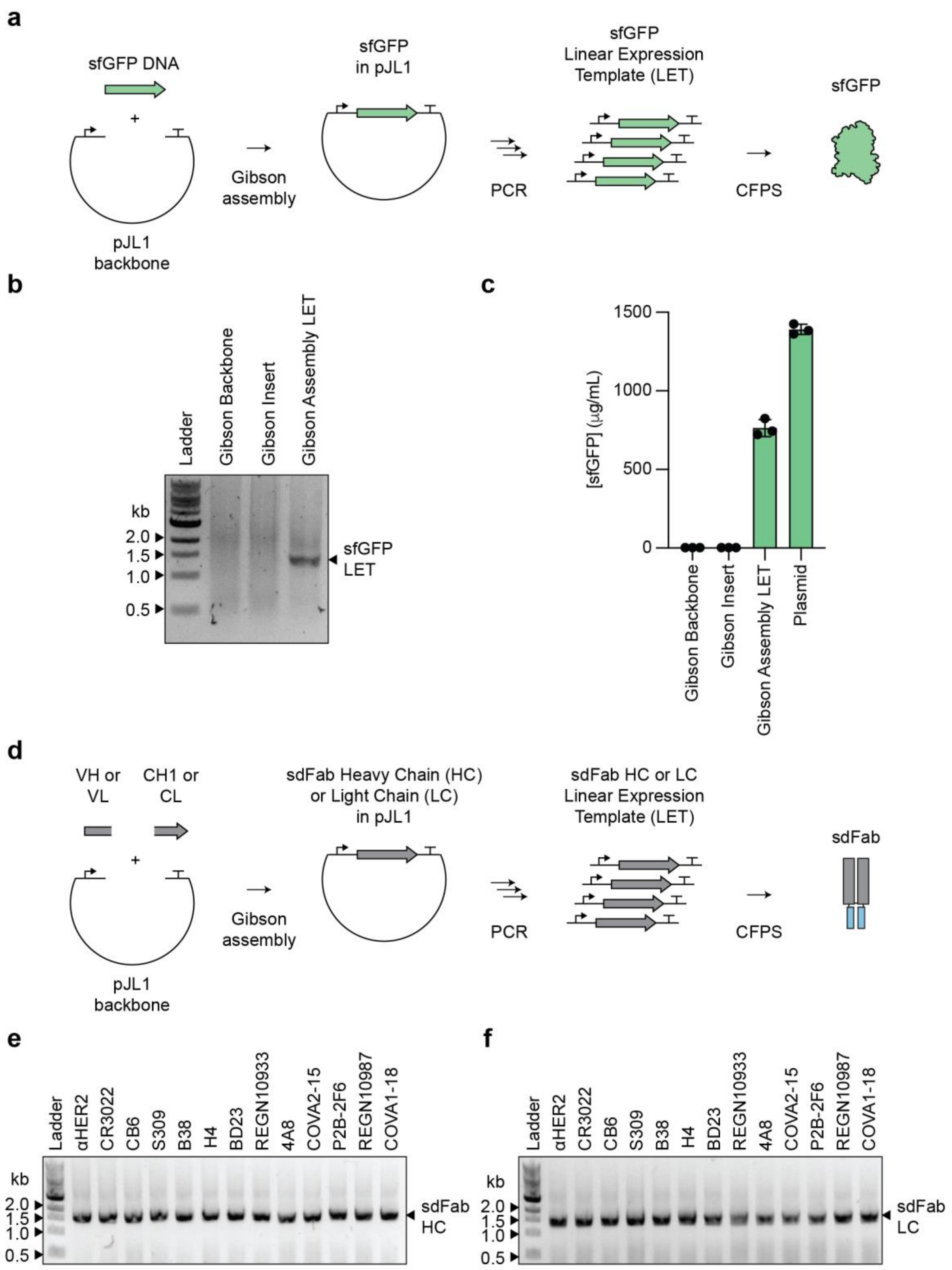


**Supplementary Figure 1 | The cell-free DNA assembly and amplification workflow**. **a**, Schematic of the cell-free DNA assembly and amplification protocol for generating sfGFP linear expression template for CFPS. **b**, Agarose gel of amplified LET PCR products of Gibson assembly reactions. Backbone only and insert only conditions included as negative controls. Labeled band indicates assembly and amplification of the correct length PCR product. **c**, sfGFP yields in Origami$^{TM}$ B(DE3) CFPS from cell-free assembled linear expression templates and purified plasmid (n = 3 independent replicates ± SEM). **d**, Schematic of the cell-free DNA assembly and amplification protocol for generating sdFab linear expression template for CFPS. **e**, Agarose gel of amplified sdFab heavy chain (HC) LET PCR products. Labeled bands indicate assembly and amplification of the correct length PCR product. **f**, Agarose gel of amplified sdFab light chain (LC) LET PCR products. Labeled bands indicate assembly and amplification of the correct length PCR product.

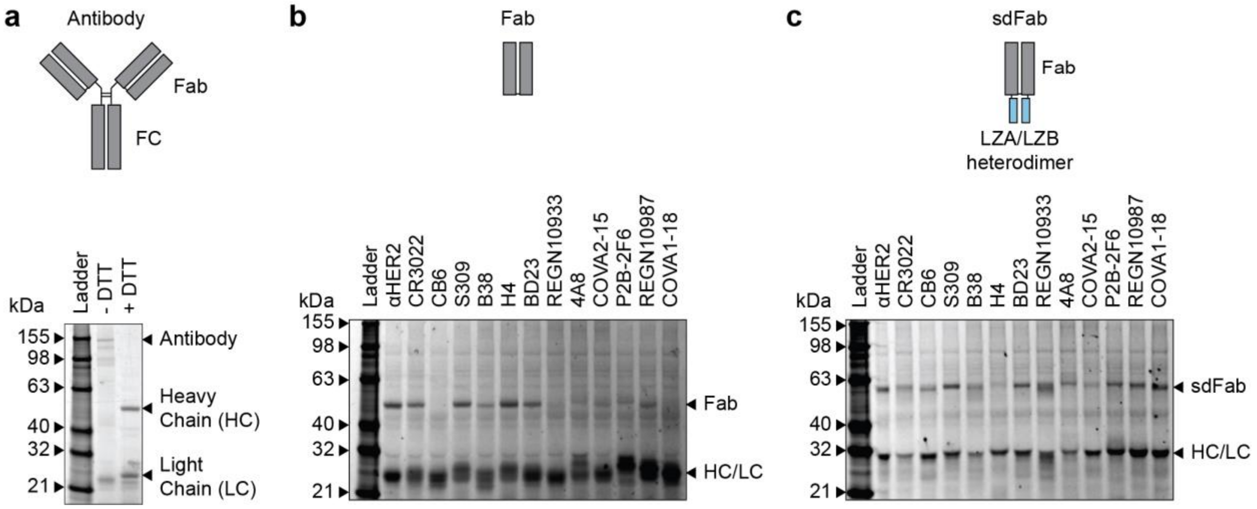


**Supplementary Figure 2 | Development of an Origami$^{TM}$ B(DE3) CFPS system for the expression of antibodies and antibody fragments**. **a-c**, SDS PAGE of antibodies and antibody fragments manufactured in CFPS. Samples were fluorescently labeled with the FluoroTect$^{TM}$ reagent during protein synthesis. **a**, Expression and assembly of full-length Trastuzumab (αHER2). The antigen-binding fragment (Fab) and fragment, crystallizeable (FC) domains are labeled. **b**, Expression and assembly of a panel of 13 Fabs. **c**, Expression and assembly of a panel of 13 sdFabs. The leucine zipper heterodimer (LZA and LZB) assisting with Fab assembly are labeled.

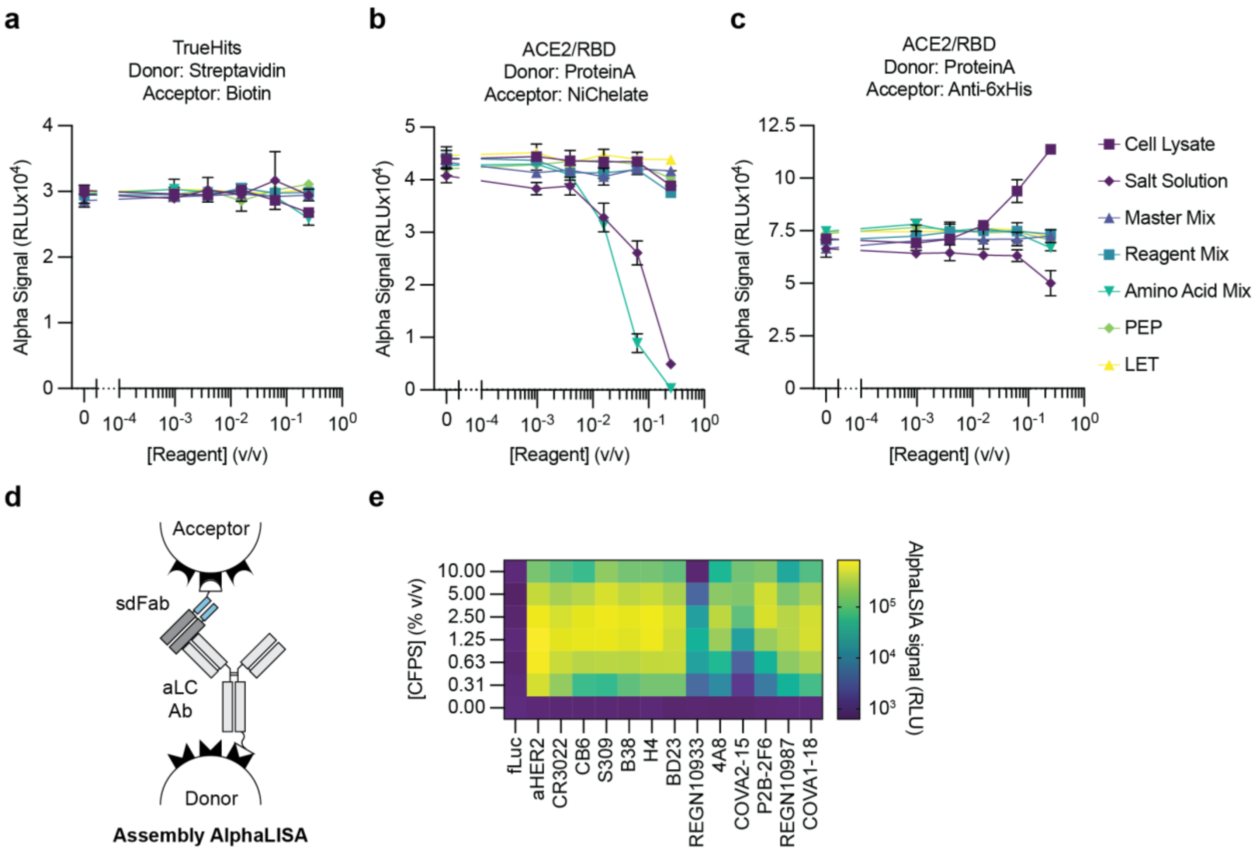


**Supplementary Figure 3 | AlphaLISA for profiling antibody protein-protein interactions in CFPS**. **a-c**, Evaluation of the effect of CFPS reagents on AlphaLISA. Concentrations are plotted as v/v fraction of the final concentration of the reagent in a CFPS reaction. Reagents were diluted in water at the concentration they normally reside at in CFPS. Reagents were tested in mixtures that were used to assemble CFPS reactions. The salt solution contains 8 mM magnesium glutamate, 10 mM ammonium glutamate, and 130 mM potassium glutamate. Master Mix contains 1.2 mM ATP, 0.85 mM GTP, 0.85 mM UTP, 0.85 mM CTP, 0.03 mg/mL folinic acid, and 0.17 *E. coli* tRNA. Reagent Mix contains 0.4 mM NAD, 0.27 mM CoA, 4 mM oxalic acid, 1 mM putrescine, 1.5 mM spermidine, and 57 mM HEPES. Amino Acid Mix contains 2 mM of all 20 amino acids. PEP is 30 µM phosphoenolpyruvate. LET is 0.066 v/v fraction unpurified PCR mix containing the LET for sfGFP. **a**, Evaluation of the effect of CFPS reagents on AlphaLISA detection chemistry using the TrueHits kit. Biotin and Streptavidin labeled beads associate directly with one another and serve as a control for reagents impacting the AlphaLISA measurement chemistry. **b**, Evaluation of the effect of CFPS reagents on AlphaLISA detection of the SARS-CoV-2 RBD and ACE2 interaction measured by the Protein A donor bead and Ni Chelate acceptor bead. The Salt Solution and Amino Acids Mix inhibit immobilization of his-tagged proteins on the NiChelate bead. **c**, Evaluation of the effect of CFPS reagents on AlphaLISA detection of the SARS-CoV-2 RBD and ACE2 interaction measured by the Protein A donor bead and anti-6xhis acceptor bead. The Salt Solution and Amino Acids Mix inhibit immobilization of his-tagged proteins on the NiChelate bead. **d**, Schematic of AlphaLISA setup for measuring sdFab assembly. **e**,

Assembly AlphaLISA measurement of a panel of sdFabs. AlphaLISA signal is indicative of sdFab assembly, though the signal is subject to the hook effect[54] resulting in a lower signal at higher concentrations.

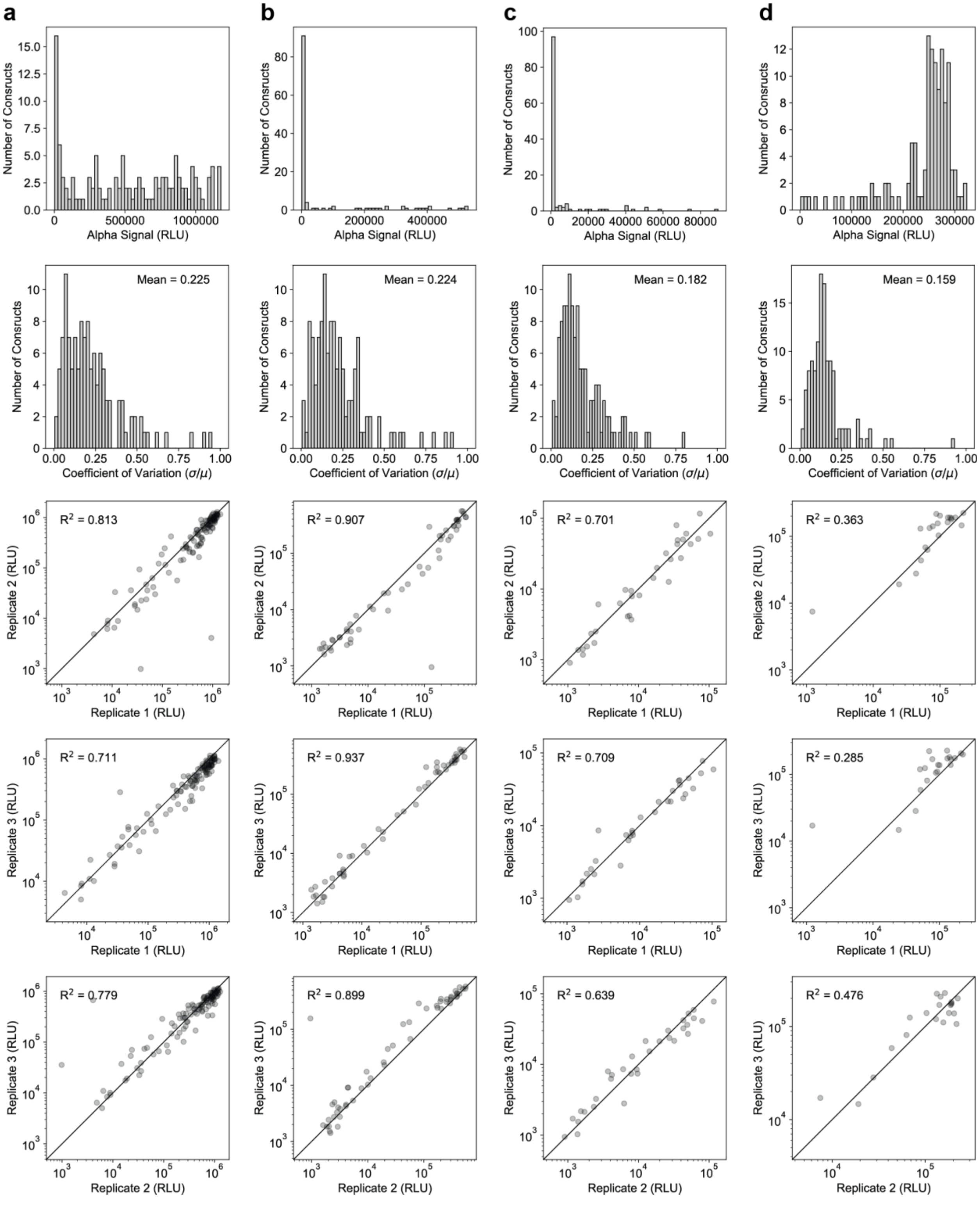


**Supplementary Figure 4 | Analysis of variability in AlphaLISA replicates**. **a-d**, From top to bottom: Histogram of raw AlphaLISA values (mean of n = 3 independent replicates) to visualize the spread of the data. A histogram of coefficient of variation (standard deviation divided by the mean) to visualize the typical error within a sample with the mean coefficient of variation displayed on the plot. Parity plots of the three replicates were fit to the line y=x to visualize the consistency of replicates with the corresponding $R^2$ value is displayed on the chart. Only values found to be significantly different from the background are plotted ($p < 0.05$, two-sided t-test adjusted for multiple comparisons using FDR with a family-wise error rate of 5%) **a**, sdFab assembly AlphaLISA. **b**, SARS-CoV-2 S trimer binding AlpahLISA. **c**, SARS-CoV-2 RBD binding AlphaLISA. **d**, sdFab competition with ACE2 for the SARS-CoV-2 RBD AlphaLISA.

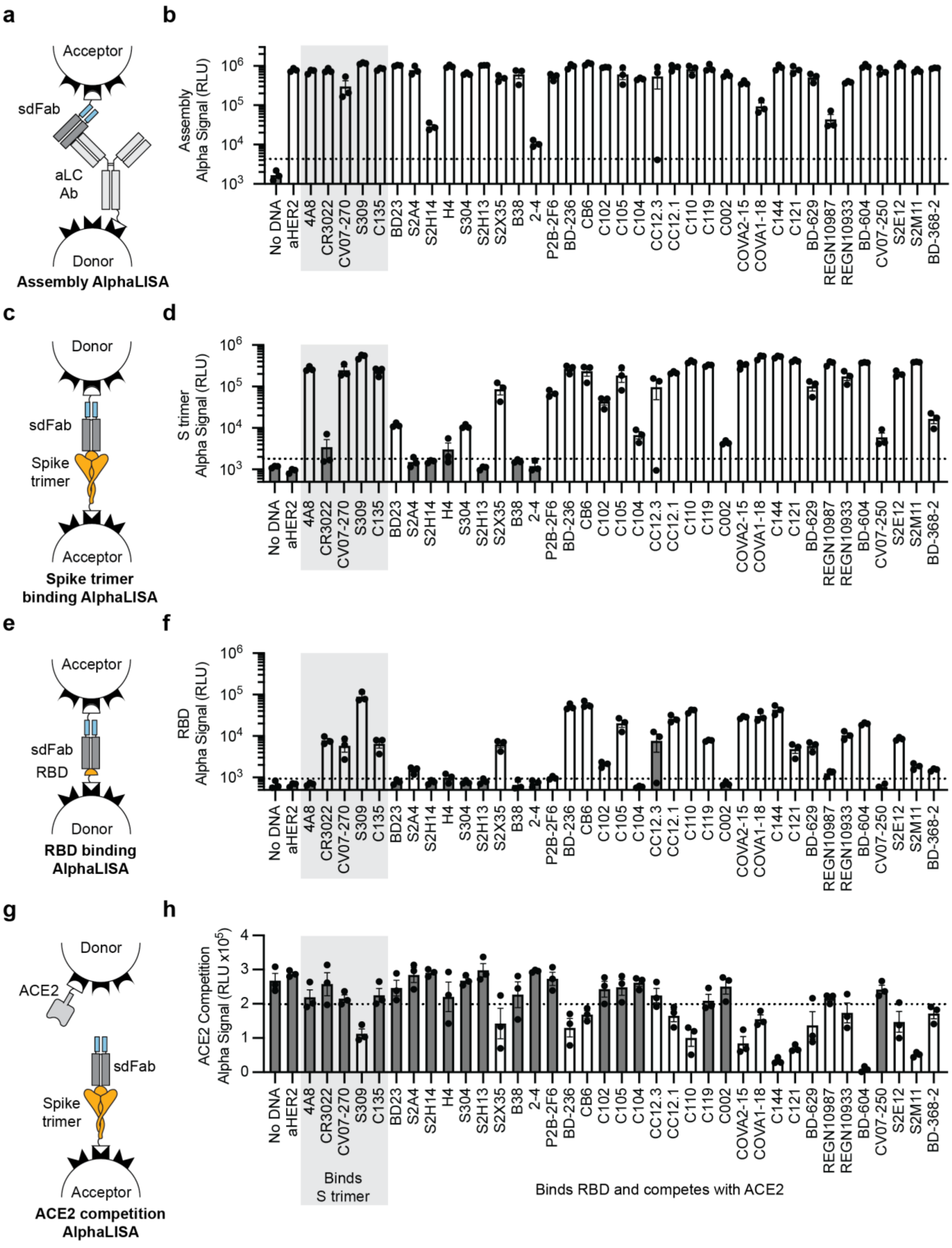
a
Acceptor
sdFab
aLC
Ab
Donor
Assembly AlphaLISA
b
Assembly
Alpha Signal (RLU)
c
Donor
sdFab
Spike
trimer
Acceptor
Spike trimer
binding AlphaLISA
d
S trimer
Alpha Signal (RLU)
e
Acceptor
sdFab
RBD
Donor
RBD binding
AlphaLISA
f
RBD
Alpha Signal (RLU)
g
Donor
ACE2
sdFab
Spike
trimer
Acceptor
ACE2 competition
AlphaLISA
h
ACE2 Competition
Alpha Signal (RLU x10$^5$)
No DNA
aHER2
4A8
CR3022
CV07-270
S309
C135
BD23
S2A4
S2H14
H4
S304
S2H13
S2X35
B38
2-4
P2B-2F6
BD-236
CB6
C102
C105
C104
CC12.3
CC12.1
C110
C119
C002
COVA2-15
COVA1-18
C144
C121
BD-629
REGN10987
REGN10933
BD-604
CV07-250
S2E12
S2M11
BD-368-2
Binds
S trimer
Binds RBD and competes with ACE2

**Supplementary Figure 5 | AlphaLISA profiling of 38 published antibodies**. Data are a more quantitative depiction of the data for 38 of the antibodies in the heatmaps in Fig. 2a-b. **a**, Schematic depicting AlphaLISA setup for measuring sdFab assembly. **b**, AlphaLISA measurement of sdFab assembly. **c**, Schematic depicting AlphaLISA setup for measuring S trimer binding. **d**, AlphaLISA measurement of sdFab binding to the SARS-CoV-2 S trimer. **e**, Schematic depicting AlphaLISA setup for measuring RBD binding. **f**, AlphaLISA measurement of sdFab binding to the SARS-CoV-2 RBD. **g**, Schematic depicting AlphaLISA setup for measuring sdFab competition with ACE2 for the RBD. **h**, AlphaLISA measurement of sdFab competition with ACE2 for the RBD. All AlphaLISA data are the mean of three independent replicates ± the SEM. The dashed line indicates three standard deviations away from the background. Samples determined not to be significantly distinguished from the background ($p>0.05$ two-sided t-test corrected using the FDR procedure) have bars that are filled dark grey. The samples are ranked within each category from worst (left) to best (right) neutralizers by their minimum neutralization $IC_{50}$ value.

| Reference | DOI | Antibodies screened | Antibodies $IC_{50}$ < 0.01 µg/mL | Antibodies $IC_{50}$ < 0.25 µg/mL |
|---|---|---|---|---|
| Kreye et al.[29] | 10.1016/j.cell.2020.09.049 | 598 | 5 | 40 |
| Cao et al.[36] | 10.1016/j.cell.2020.05.025 | 216 | 1 | 22 |
| Wu et al.[35] | 10.1126/science.abc2241 | 17 | 0 | 1 |
| Liu et al.[32] | 10.1038/s41586-020-2571-7 | 252 | 9 | NR |
| Ju et al.[30] | 10.1038/s41586-020-2380-z | 206 | 0 | 3 |
| Brouwer et al.[25] | 10.1126/science.abc5902 | 84 | 2 | 9 |
| Hansen et al.[38] | 10.1126/science.abd0827 | "Thousands" | 6 | NR |
| Robbani et al.[55] | 10.1038/s41586-020-2456-9 | 94 | 9 | 43 |

**Supplementary Table 1 | Summary of antibody screening studies designed to identify SARS-CoV-2 neutralizing antibodies.** Studies were evaluated for their efficiency at identifying potent neutralizing antibodies. For each study, the total number of antibodies evaluated as well as the number of neutralizing antibodies with a neutralization $IC_{50}$ less than 0.01 µg/mL and 0.25 µg/mL were summarized. Either authentic- or pseudovirus neutralization $IC_{50}$ was considered based on the breadth of the reported data. The 0.01 µg/mL cutoff was chosen because this value is approximately within an order of magnitude of the most potent reported neutralizing antibodies[32]. 0.25 µg/mL was chosen as a practical cutoff for moderately potent neutralizing antibodies[29]. For Hansen *et al.*[38] the neutralization potencies were reported in M and were converted to µg/mL assuming an antibody molecular weight of 150 kDa. For this analysis, only the studies containing antibodies expressed in this manuscript were used. Furthermore, only studies whose purpose was to identify neutralizing antibodies from a large set of candidates were considered. Studies that didn’t describe the results of their antibody discovery process in sufficient detail to collect the desired information were omitted. NR indicates not reported in sufficient detail to determine.

**Supplementary Table 2 | Reported parameters and expected behaviors for tested antibodies.** Information about antibody target epitope, pseudo- or authentic virus neutralization $IC_{50}$, and equilibrium dissociation constant  from literature about the antibodies used in this study are summarized. Data are presented in two tabs, one for antibodies from diverse sources and one for the Brouwer *et al.* data set, in a separate .xlsx file.

**Supplementary Table 3 | Variable heavy and light chain sequences of tested antibodies.** Sequences are classified by their heavy (VH) or light chain (VL) as well as the light chain class (kappa or lambda). The variable domain protein sequence, the variable domain *E. coli* codon-optimized DNA sequence, and the ordered DNA sequence containing all additional (Gibson assembly homology, n-terminal expression tag, etc.) sequences are listed. Note that the antibodies COVA2-15 and COVA1-18 were ordered and evaluated twice in this data set, and there are thus two separate entries for the two sets of sequences. Sequences are listed in a single tab in a separate .xlsx file.

**Supplementary Table 4 | Summary of observed antibody binding significance statistics and comparison to literature.** Results from the AlphaLISA screen are compared against literature data and a qualitative match with the reported values are summarized. Comments on the possible origin of observed inconsistent results are listed for all data with reported neutralization IC50 values less than 10 μg/mL. Data are presented in two tabs, one for antibodies from diverse sources and one for the Brouwer *et al.* data set, in a separate .xlsx file.

**Supplementary Table 5 | Raw and processed data for AlphaLISA measurements for the 120 antibodies evaluated in this manuscript.** The individual replicates, average, standard deviation, coefficient of variation, p-value from a two-sided t-test against the background, and the FDC corrected p-value are reported. Note that the antibodies COVA2-15 and COVA1-18 were ordered and evaluated twice in this data set, and there are thus two separate entries for the two sets of sequences. Data are presented in four tabs, one for each AlphaLISA measurement modality, in a separate .xlsx file.

### Materials and Methods

#### Antibody sdFab sequence design

sdFabs were assembled based on a modified version of previously published protocols[17–19]. Antibody sequences were collected from literature and their light chains were classified as either kappa or lambda via the terminal residue of the J-segment in the VL domain. The VH and VL domains were subsequently fused to their corresponding human constant heavy (Uniprot P0DOX5) or human constant light (kappa CL Uniprot P01834 or lambda 1 CL Uniprot P0CG04) chains. At the N-terminus of the VH and VL domains, we chose to include a modified expression tag based on the first 5-residues of the *E. coli* chloramphenicol acetyltransferase gene followed by a Tobacco Etch Virus (TEV) protease cleavage site (protein sequence: MEKKIENLYFQS, DNA sequence: atggagaaaaaaatcgaaaacctgtacttccagagc)[56] as opposed to the previously published SKIK tag[57]. The heavy chain was fused to the LZA heterodimer subunit (AQLEKELQALEKENAQLEWELQALEKELAQK) and a strep II tag. The light chain was fused to the LZB heterodimer subunit (AQLKKKLQALKKKNAQLKWKLQALKKKLAQK). Examples of the three types of antibody sequences are detailed below, with the important sequence features highlighted in square brackets [].

sdFab heavy chain constant strepII tagged:
[MEKKIENLYFQS][VH_Sequence][ASTKGPSVFPLAPSSKSTSGGTAALGCLVKDYFPE
PVTVSWNSGALTSGVHTFPAVLQSSGLYSLSSVVTVPSSSLGTQTYICNVNHKPSNTK
VDKKVEPKSC]GGGGS[AQLEKELQALEKENAQLEWELQALEKELAQK]GSSA[WSHP
QFEK]

sdFab light chain kappa:
[MEKKIENLYFQS][VL_Sequence][RTVAAPSVFIFPPSDEQLKSGTASVVCLLNNFYPR
EAKVQWKVDNALQSGNSQESVTEQDSKDSTYSLSSTLTLSKADYEKHKVYACEVTHQ
GLSSPVTKSFNRGEC]GGGGS[AQLKKKLQALKKKNAQLKWKLQALKKKLAQK]

sdFab light chain lambda 1:
[MEKKIENLYFQS][VL_Sequence][GQPKANPTVTLFPPSSEELQANKATLVCLISDFYP
GAVTVAWKADGSPVKAGVETTKPSKQSNNKYAASSYLSLTPEQWKSHRSYSCQVTH
EGSTVEKTVAPTECS]GGGGS[AQLKKKLQALKKKNAQLKWKLQALKKKLAQK]

#### DNA assembly and linear expression template (LET) generation

Proteins to be manufactured via CFPS were codon-optimized using the IDT codon optimization tool and ordered as double-stranded linear DNA containing the desired Gibson assembly overhangs from IDT or GenScript. sfGFP was ordered containing the two pJL1 Gibson assembly overhangs. Antibody VH DNA was ordered with the pJL1 5' and the human IgG1 heavy chain constant 5' Gibson overhangs. Antibody VL DNA was ordered with the pJL1 5' and the human Ig light chain kappa or lambda 1 Gibson assembly overhangs. DNA was resuspended at a concentration of 50 ng/μL and used without amplification.

Additional linear DNA components for Gibson assembly (pJL1 backbone, sdFab heavy chain constant strepII tagged, sdFab light chain kappa constant, sdFab light chain lambda 1 constant) were ordered as gblocks from IDT. These components were amplified using PCR using Q5 Hot Start DNA polymerase (NEB, M0493L) following manufacturer instructions. Amplified DNA was purified using the DNA Clean and Concentrate Kit (Zymo Research, D4006) and diluted to a concentration of 50 ng/μL. Sequences of the utilized components are listed below, with Gibson assembly sequences being denoted by underlined lowercase text and primers for a given amplicon being listed below the DNA sequence.

Gibson assembly overhangs:
pJL1 5' Gibson: tttgtttaactttaagaaggagatatacat
pJL1 3' Gibson: gtcgaccggctgctaacaaagcccgaaagg
Human IgG1 heavy chain constant 5' Gibson: gcgtcaacaaaaggtccttcagttttcccattagcccct
Human Ig light chain kappa 5' Gibson: cgcacggtcgcggcgccgtctgtctttattttcctcct
Human Ig light chain lambda 5' Gibson: ggccaacccaaagcaaacccaactgtcactttgttcccg

Linear pJL1 plasmid backbone (Addgene plasmid # 69496):
gtcgaccggctgctaacaaagcccgaaaggAAGCTGAGTTGGCTGCTGCCACCGCTGAGCAAT
AACTAGCATAACCCCTTGGGGCCTCTAAACGGGTCTTGAGGGGTTTTTTGCTGAAA
GCCAATTCTGATTAGAAAAACTCATCGAGCATCAAATGAAACTGCAATTTATTCATA
TCAGGATTATCAATACCATATTTTTGAAAAAGCCGTTTCTGTAATGAAGGAGAAAAC
TCACCGAGGCAGTTCCATAGGATGGCAAGATCCTGGTATCGGTCTGCGATTCCGA
CTCGTCCAACATCAATACAACCTATTAATTTCCCCTCGTCAAAAATAAGGTTATCAA
GTGAGAAATCACCATGAGTGACGACTGAATCCGGTGAGAATGGCAAAAGCTTATG
CATTTCTTTCCAGACTTGTTCAACAGGCCAGCCATTACGCTCGTCATCAAAATCACT
CGCATCAACCAAACCGTTATTCATTCGTGATTGCGCCTGAGCGAGACGAAATACGC
GATCGCTGTTAAAAGGACAATTACAAACAGGAATCGAATGCAACCGGCGCAGGAA
CACTGCCAGCGCATCAACAATATTTTCACCTGAATCAGGATATTCTTCTAATACCTG
GAATGCTGTTTTCCCGGGGATCGCAGTGGTGAGTAACCATGCATCATCAGGAGTA
CGGATAAAATGCTTGATGGTCGGAAGAGGCATAAATTCCGTCAGCCAGTTTAGTCT
GACCATCTCATCTGTAACATCATTGGCAACGCTACCTTTGCCATGTTTCAGAAACAA
CTCTGGCGCATCGGGCTTCCCATACAATCGATAGATTGTCGCACCTGATTGCCCGA
CATTATCGCGAGCCCATTTATACCCATATAAATCAGCATCCATGTTGGAATTTAATC
GCGGCTTCGAGCAAGACGTTTCCCGTTGAATATGGCTCATAACACCCCTTGTATTA
CTGTTTATGTAAGCAGACAGTTTTATTGTTCATGATGATATATTTTTATCTTGTGCAA
TGTAACATCAGAGATTTTGAGACACAACGTGAGATCAAAGGATCTTCTTGAGATCC
TTTTTTTCTGCGCGTAATCTGCTGCTTGCAAACAAAAAAACCACCGCTACCAGCGG
TGGTTTGTTTGCCGGATCAAGAGCTACCAACTCTTTTTCCGAAGGTAACTGGCTTC
AGCAGAGCGCAGATACCAAATACTGTTCTTCTAGTGTAGCCGTAGTTAGGCCACCA
CTTCAAGAACTCTGTAGCACCGCCTACATACCTCGCTCTGCTAATCCTGTTACCAG
TGGCTGCTGCCAGTGGCGATAAGTCGTGTCTTACCGGGTTGGACTCAAGACGATA
GTTACCGGATAAGGCGCAGCGGTCGGGCTGAACGGGGGGTTCGTGCACACAGCC
CAGCTTGGAGCGAACGACCTACACCGAACTGAGATACCTACAGCGTGAGCTATGA
GAAAGCGCCACGCTTCCCGAAGGGAGAAAGGCGGACAGGTATCCGGTAAGCGGC
AGGGTCGGAACAGGAGAGCGCACGAGGGAGCTTCCAGGGGGAAACGCCTGGTAT

CTTTATAGTCCTGTCGGGTTTCGCCACCTCTGACTTGAGCGTCGATTTTTGTGATG
CTCGTCAGGGGGGCGGAGCCTATGGAAAAACGCCAGCAACGCGATCCCGCGAAA
TTAATACGACTCACTATAGGGAGACCACAACGGTTTCCCTCTAGAAATAATtttgtttaact
ttaagaaggagatatacat

pJL1_F gtcgaccggctgcta
pJL1_R atgtatatctccttcttaaagttaaacaaaattatttcta

Linear sdFab heavy chain constant strepII tagged:
gcgtcaacaaaaggtccttcagttttcccattagcccctTCTTCTAAGTCAACTAGTGGCGGTACTGCC
GCTCTTGGGTGTTTGGTTAAAGATTACTTCCCAGAACCGGTTACGGTCTCGTGGAA
CTCTGGTGCACTGACATCGGGCGTACATACATTTCCCGCAGTTTTGCAGTCTTCGG
GACTGTATTCTCTTTCATCGGTGGTTACAGTCCCTAGCTCTTCCCTGGGTACACAG
ACCTACATTTGTAATGTTAATCATAAGCCGAGTAATACTAAGGTGGATAAAAAGGTG
GAACCGAAGTCTTGTGGTGGTGGCGGGTCAGCTCAACTGGAGAAGGAGTTACAG
GCACTGGAAAAAGAGAATGCTCAACTTGAGTGGGAATTACAGGCGTTAGAAAAAGA
ACTGGCCCAGAAGGGTTCTAGCGCATGGTCACATCCCCAGTTCGAAAAATAAgtcga
ccggctgctaacaaagcccgaaagg

IgGC_F: GCGTCAACAAAAGGTCCTTCAGTTTTC
pJL1_3'Gib_R: CCTTTCGGGCTTTGTTAGCAGC

Linear sdFab light chain kappa constant:
cgcacggtcgcggcgccgtctgtctttatttttcctcctTCTGATGAACAGCTTAAATCTGGGACAGCTT
CTGTTGTATGTTTATTAAACAACTTTTACCCGCGTGAGGCAAAAGTTCAATGGAAGG
TAGACAACGCACTGCAAAGCGGAAATTCGCAGGAGTCAGTTACCGAACAGGATTC
CAAGGATAGTACCTACTCCTTAAGTTCAACATTAACCCTGTCAAAGGCGGACTATG
AAAAACATAAGGTATATGCCTGCGAAGTAACTCATCAGGGCTTATCATCCCCAGTT
ACAAAATCTTTCAACCGTGGAGAATGCGGCGGCGGAGGTAGCGCGCAGCTTAAGA
AAAAATTGCAAGCCCTTAAAAAAAAAAATGCCCAACTTAAATGGAAGCTGCAAGCC
TTAAAAAAGAAATTGGCGCAGAAGTAAgtcgaccggctgctaacaaagcccgaaagg

kLC_F: TCGCGGCGCCGTCTG
pJL1_3'Gib_R: CCTTTCGGGCTTTGTTAGCAGC

Linear sdFab light chain lambda 1 constant:
ggccaacccaaagcaaacccaactgtcactttgttcccgCCCTCAAGCGAGGAACTTCAGGCTAATA
AGGCCACGCTTGTTTGCCTGATCTCAGACTTTTATCCCGGTGCCGTAACAGTGGCT
TGGAAGGCAGATGGTTCGCCGGTCAAAGCGGGCGTGGAAACTACAAAGCCATCG
AAACAGTCAAACAATAAATATGCGGCATCAAGTTACTTGAGCCTTACCCCAGAACA
GTGGAAGTCACACCGCTCGTACAGTTGTCAAGTTACACACGAGGGAAGTACAGTT
GAAAAGACCGTTGCCCCAACTGAATGTTCAGGCGGTGGTGGCTCAGCGCAGTTAA
AGAAAAAACTGCAGGCTTTGAAGAAAAAGAATGCTCAATTAAAGTGGAAATTGCAG
GCGTTGAAGAAGAAACTTGCGCAGAAGTAAgtcgaccggctgctaacaaagcccgaaagg

lLC_F: GGCCAACCCAAAGCAAACC

pJL1_3'Gib_R: CCTTTCGGGCTTTGTTAGCAGC

Gibson assembly was used to assemble protein open reading frame DNA with the pJL1 backbone following the published protocol with the addition of 3.125 µg/mL of ET SSB (NEB, product no. M2401S)[58,59]. 20 ng of purified, linear pJL1 backbone, 20 ng of purified, linear sdFab VH or VL constant DNA, and 20 ng of the protein open reading frame insert were combined in 2 µL Gibson assembly reactions and incubated at 50°C for 30 minutes. The unpurified assembly reactions were diluted in 40 µL of nuclease-free water (Fisher Scientific, AM9937) and 1 µL of the diluted reaction was used as the template for a PCR to generate linear expression templates (LETs) for CFPS. Linear expression templates were amplified via PCR using the pJL1_LET_F (ctgagatacctacagcgtgagc) and pJL1_LET_R (cgtcactcatggtgatttctcacttg) primers in a 50 µL PCR reaction using the Q5 Hot Start DNA polymerase (NEB, M0493L) following manufacturer instructions.

**Additional Protein DNA Sequences**

The DNA sequence of the *P. pyralis* luciferase containing a c-terminal strepII tag (fLuc, Uniprot Q27758) used as a negative control is below and was cloned into the pJL1 vector.

atggaagacgctaagaacattaagaagggacctgctccattctacccctcgaagacggcactgcaggtgagcagcttc
ataaagcgatgaagcgttatgcgttagttcctggcacgatcgccttcactgacgcgcacatcgaagtcaatatcacctacg
ctgaatactttgagatgagtgtgcgtctggcggaagccatgaagcgttatggccttaacacgaaccaccgcatcgttgtttgt
agcgagaattccttacaattcttcatgcccgtccttggcgcgctgtttattggtgtggccgttgcaccagccaatgacatctata
atgagcgcgagttgttgaactccatgaacatttctcaaccaacagtggtgttcgtttcaaagaaaggcttacagaaaatctta
aacgttcaaaagaaactgccgattatccagaagatcatcattatggatagtaagactgactaccagggcttccagtcaatg
tatacattcgtgacgagtcacctgcccccgggttttaacgagtacgactttgtcccagagagctttgatcgcgacaagacca
tcgccctcattatgaatagcagtggttcgacgggtagcccaaagggagtggccctgccccatcgtaccgcgtgcgtccgttt
ctcccatgcccgcgacccaattttcggcaatcaaatcatccccgacacggcaatcttgtcggtcgtcccgtttcaccatggct
ttggaatgtttacgacactcggttacctcatctgcggtttccgcgtcgttctgatgtatcgcttcgaggaagagttgttcttacgttc
gcttcaggactacaagattcaatccgcccttctggtccccactttgttcagtttctttgctaagagcaccttaattgataagtatg
acctctccaacttacacgagattgcgagcggtggtgctcccctcagcaaagaggttggagaggcggttgctaagcgttttc
atctgcccggtatccgtcaaggttacggcctcaccgaaaccacttctgccattcttatcactccggaaggtgacgataagcc
tggggcagtgggtaaagttgtacccttcttcgaggctaaggttgtggatttagatacggggaagaccttaggtgtgaaccag
cgcggtgaactgtgcgttcgcggtccgatgattatgtcgggttatgttaatgaccccgaggctacgaacgcgcttatcgataa
ggacggttggcttcattccggcgacatcgcttactgggatgaggatgagcacttcttcatcgttgaccgtctgaagagtctcat
caagtataagggatgtcaagtcgctccggcagagttagagagcatcttactccagcaccctaatatcttcgatgctggggtt
gccgggctcccaggcgacgatgccggcgagctgccggcggtagttgttttagagcatggcaagaccatgaccgaa
aaggagattgtagactacgtcgcgagtcaagtaaccacagcgaagaagctccgcggtggagtggtctttgttgacgaggt
gcctaaaggcctgacgggcaaacttgacgcgcgtaagatccgtgagatcctcatcaaagcgaagaagggtgggaaga
gtaagctggggagttcaggttggtcccacccgcaatttgagaagtga

**Cell extract preparation for cell-free protein synthesis**

E. coli Origami™ B(DE3) (Novagen, 70837) extracts were prepared using a modified version of established protocols[60,61]. Briefly, a 150 mL Origami™ B(DE3) starter culture was inoculated in LB from a glycerol stock and cultured in a 250 mL baffled flask at 37 °C for 16 hours. The 2xYTP was prepared lacking glucose in 75% of the final volume and sterilized using an autoclave. A 4x glucose solution was prepared and autoclaved separately, then added to the medium immediately before use. The starter cultures were used to inoculate 1 L of 2xYTPG media (16 g/L tryptone, 10 g/L yeast extract, 5 g/L sodium chloride, 7 g/L potassium phosphate dibasic, 3 g/L potassium phosphate monobasic, 18 g/L glucose) in a 2.5 L Full-Baffle Tunair shake flask at an initial OD600 of 0.08. Cells were cultured at 37 °C at 220 RPM in a shaking incubator. Cultures were grown until OD600 0.4-0.6, at which point the expression of T7 RNA polymerase was induced by the addition of IPTG to a final concentration of 0.5 mM. Cells were harvested at an OD600 of 2.5 via centrifugation at 12,000g for 1 minute at 4 °C. Cell pellets were washed three times with 25 mL S30 buffer per 50 mL culture (10 mM Tris Acetate pH 8.2, 14 mM Magnesium Acetate, and 60 mM Potassium Acetate). Pellets were resuspended in 1 mL S30 buffer per gram of cell mass. Cell suspensions were lysed using a single pass on an Avestin EmulsiFlex-B15 Homogenizer at a lysis pressure of 24,000 PSI. Cell debris was separated via centrifugation at 18,000g for 20 minutes, and the clarified lysate was collected, flash-frozen in liquid nitrogen, and stored at -80 °C.

**DsbC and FkpA expression and purification**
Protein expression, purification, and his tag removal were performed similarly to previously reported[62]. DsbC (Uniprot P0AEG6, residues 21-236) and FkpA (Uniprot P45523, residues 26-270) were ordered as gBlocks from IDT containing a c-terminal, TEV cleavable his tag (GSENLYFQSGSHHHHHHHHHH) and cloned into pET28a. Plasmids were transformed into BL21 Star™ DE3, plated on LB agar, and cultured overnight at 37°C. 1 L of Overnight Express TB (Fisher Scientific, 71491-4) was inoculated by scraping all colonies on a transformation plate and cultured at 37°C in 2.5 L tunair flasks (IBI Scientific, SS-8003) at 220 rpm overnight. Cells were harvested, resuspended at a ratio of 1 g cell mass to 4 mL resuspension buffer (50 mM HEPES pH 7.5, 500 mM NaCl, 1X HALT protease inhibitor without EDTA (Fisher Scientific, 78429), 1 mg/mL lysozyme, 62.5 U/mL cell suspension of benzonase (Sigma Aldrich, E1014-25KU)) and lysed using an Avestin B15 homogenizer at 24,000 PSI. The lysate was spun down 14,000 x g for 10 min and the clarified supernatant was incubated with Ni-NTA Agarose (Qiagen, 30230) for 60 min on an end-over-end shaker. The resin was spun down 2,500 x g for 2 min, the supernatant removed, resuspended in wash buffer (50 mM HEPES pH 7.5, 500 mM NaCl, 50 mM Imidazole), loaded on a gravity flow column, and subsequently washed with 20X resin volumes of wash buffer. Protein was eluted using elution buffer (50 mM HEPES pH 7.5, 500 mM NaCl, 500 mM Imidazole) and exchanged into 50 mM HEPES pH 7.4, 150 mM NaCl using PD-10 desalting columns (Cytvia, 17-0851-01) according to manufacturer instructions.

His tags were removed via cleavage by ProTEV Plus (Promega, V6102). Before cleavage, 10% v/v glycerol was added to the protein. ProTEV Plus was added to a concentration of 0.5 U/µg purified protein and DTT was added to a concentration of 1 mM. Cleavage reactions were carried out at 30°C for 4 h. Free His tag and ProTEV Plus were removed by incubating with Ni-NTA Agarose for 1 hour at 4°C and collecting the supernatant. Proteins were subsequently concentrated to > 1mg/mL (Millipore, UFC800396). His tag removal was validated via SDS-PAGE and the AlphaScreen Histidine (Nickel Chelate) Detection Kit (Perkin Elmer, 6760619C).

**Cell-free protein synthesis**

CFPS reactions were composed of the following reagents: 8 mM magnesium glutamate, 10 mM ammonium glutamate, 130 mM potassium glutamate, 1.2 mM ATP, 0.5 mM of each CTP, GTP, and UTP. 0.03 mg/mL folinic acid, 0.17 mg/mL *E. coli* MRE600 tRNA (Roche 10109541001), 100 mM NAD, 50 mM CoA, 4 mM oxalic acid, 1 mM putrescine, 1 mM spermidine, 57 mM HEPES pH 7.2, 2 mM of each amino acid, 33.3 mM PEP, 20% v/v *E. coli* extract, varying concentrations of DNA template, and the remainder water. The preparation of these reagents has been described in detail elsewhere[63]. For DNA templates, plasmids were used at a concentration of 8 nM, and unpurified linear PCR products were used at 6.66 %v/v. For the expression of antibodies, each template was added to a final concentration of 6.66 %v/v. For antibody and sdFab expression 4 mM oxidized glutathione, 1 mM reduced glutathione, 14 µM of purified DsbC, and 50 µM FkpA were also supplemented to the reactions. Additionally, for oxidizing CFPS reactions, cell-extracts were treated with 50 mM iodoacetamide (IAM) at room temperature for 30 minutes before use in CFPS[64]. All reaction components were assembled on ice and were either run as 12 µL reactions in 1.5 mL microtubes or 2 µL reactions in 384 well plates (BioRad, HSP3801). For 2 µL reactions, components were transferred to the plate using an Echo 525 acoustic liquid handler. A mix containing all the CFPS components except for the DNA was dispensed from 384PP Plus plates (Labcyte, PPL-0200) using the BP setting. The DNA (unpurified PCR products) was dispensed from a 384LDV Plus plate (Labcyte, LPL-0200) using the GP setting. Reactions were allowed to proceed at 30 °C for 20 hours.

To quantify sfGFP fluorescence, a standard curve was prepared using previously reported methods[61]. Radioactive leucine was added to CFPS at a final concentration of 10 µM of L-[14C(U)]-leucine (Perkin Elmer NEC279E250UC, 11.1GBq/mMole), followed by precipitation of the expressed proteins and scintillation counting[65]. To quantify sfGFP fluorescence, 2 µL of a CFPS reaction was diluted in 48 µL of water in a Black Costar 96 Well Half Area Plate. Fluorescence was measured using a BioTek Synergy™ H1 plate reader with excitation and emission wavelengths of 485 and 528 respectively. Scintillation counts and fluorescence were fit to determine a standard curve for use with non-radioactive samples.

To visualize antibody assembly, proteins were labeled during CFPS with FluoroTect™ (Promega, L5001). FluoroTect™ was included in the CFPS reaction at 3.33 %v/v. After protein synthesis, RNAseA (Omega Bio-Tek, AC118) was added to 0.1 mg/mL and the sample was incubated at 37 °C for 10 minutes. Samples were subsequently denatured at 70 °C for 3 minutes, then separated via SDS-PAGE and imaged using a LI-COR Odyssey Fc imager on the 600 channel.

**AlphaLISA reactions**

AlphaLISA reactions were carried out in 50 mM HEPES pH 7.4, 150 mM NaCl, 1 mg/mL BSA, and 0.015% v/v TritonX-100 (hereafter referred to as Alpha buffer). All components were dispensed using an Echo 525 liquid handler from a 384-Well Polypropylene 2.0 Plus microplate (Labcyte, PPL-0200) using the 384PP_Plus_GPSA fluid type. All components of the AlphaLISA reactions were prepared as 4x stocks and added as 0.5 µL to the final

2 µL reaction to achieve the desired concentration. All AlphaLISA reactions were performed with CFPS reactions diluted to a final concentration of 2.5 %v/v. AlphaLISA beads were combined to prepare a 4X stock in Alpha buffer immediately before use and added to the proteins to yield a concentration of 0.08 mg/mL donor beads and 0.02 mg/mL acceptor beads in the final reaction. All reactions were incubated with AlphaLISA beads for 1 hour before measurement. AlphaLISA measurements were taken on a Tecan Infinite M1000 Pro plate reader using the AlphaLISA filter with an excitation time of 100 ms, an integration time of 300 ms, and a settling time of 20 ms. Before measurement, plates were allowed to equilibrate inside the instrument for 10 min. For measurements involving sdFabs, protein A AlphaLISA beads were avoided due to the ability of protein A to bind human subgroup VH3 Fabs[66].

The impact of CFPS reagents on AlphaLISA was determined by serially diluting the specified reagents in Alpha buffer and combining them with the specified AlphaLISA conditions. The TrueHits kit (Perkin Elmer, AL900) was used to assess the impact of the CFPS reagents on the Alpha detection chemistry. CFPS reagents were mixed with the donor and acceptor beads and incubated for 2 hours before measurement. His tagged RBD (Sino Biological, 40592-V08H) and human FC tagged human ACE2 (GenScript, Z03484) were used to evaluate the impact of CFPS reagents on capture chemistries. RBD and ACE2 were diluted in Alpha buffer, mixed at a final reaction concentration of 10 nM each, combined with the CFPS reagents, and allowed to incubate for 1 hour. Donor and acceptor beads were subsequently added and allowed to incubate for a further hour before measurement. Protein A Alpha donor beads (Perkin Elmer, AS102), Ni-Chelate AlphaLISA acceptor beads (Perkin Elmer, AL108), and anti-6xhis AlphaLISA acceptor beads (Perkin Elmer, AL178) were utilized for detection.

The commercial neutralizing antibody ACE2 competition experiment was performed with the following antibodies: nAb1 (Acro Biosystems, SAD-S35), nAb2 (Sino Biological, 40592-MM57), nAb3 (Sino Biological, 40591-MM43), nAb4 (Sino Biological, 40592-R001). ELISA $IC_{50}$ values were recorded from the product page at the time of purchase and converted to µg/mL assuming a MW of 150,000 Da if reported in M. Antibodies were serially diluted in Alpha buffer and mixed with SARS-CoV-2 RBD (Sino Biological, 40592-V02H) at a concentration of 10 nM in the final reaction and incubated for 1 hour. Mouse FC tagged human ACE2 (Sino Biological, 10108-H05H) was subsequently added and incubated for 1 hour, followed by simultaneous addition of the acceptor and donor beads. AlphaLISA detection was performed using Anti-Mouse IgG Alpha Donor beads (PerkinElmer, AS104) and Strep-Tactin AlphaLISA Acceptor beads (PerkinElmer, AL136). $IC_{50}$ values were calculated using Prism 9 by fitting the normalized data to [Inhibitor] vs. response -- Variable slope (four parameters) fit with the max constrained to a value of 1.

Assembly AlphaLISA reactions were composed of a final concentration of 10 nM of either Rabbit Anti-Human kappa light chain antibody (abcam, ab125919) or Rabbit Anti-Human lambda light chain (abcam, ab124719). AlphaLISA detection was performed using Anti-Rabbit IgG Alpha Donor beads (PerkinElmer, AS105) and Strep-Tactin AlphaLISA Acceptor beads (PerkinElmer, AL136). CFPS reaction containing the expressed sdFab of interest was mixed with the appropriate anti-light chain antibody and allowed to equilibrate for two hours before the simultaneous addition of the acceptor and donor beads.

SARS-CoV-2 S trimer binding AlphaLISA reactions were composed of a final concentration of 5 nM His-tagged SARS-CoV-2 S trimer (Acro Biosystems, SPN-C52H9). AlphaLISA detection was performed using Strep-Tactin Alpha Donor beads (PerkinElmer, AS106) and Anti-6xHis AlphaLISA Acceptor beads (PerkinElmer, AL178). CFPS reaction containing the expressed sdFab of interest was mixed with the S trimer and allowed to equilibrate for two hours before the simultaneous addition of the acceptor and donor beads.

SARS-CoV-2 RBD binding AlphaLISA reactions were composed of a final concentration of 5 nM Human Fc-tagged SARS-CoV-2 RBD (Sino Biological, 40592-V02H). AlphaLISA detection was performed using Anti-Human IgG Alpha Donor beads (PerkinElmer, AS114) and Strep-Tactin AlphaLISA Acceptor beads (PerkinElmer, AL136). CFPS reaction containing the expressed sdFab of interest was mixed with the RBD and allowed to equilibrate for two hours before the simultaneous addition of the acceptor and donor beads.

ACE2 and RBD competition AlphaLISA reactions were composed of a final concentration of 2 nM Biotinylated SARS-CoV-2 S trimer (Acro Biosystems, SPN-C82E9) and 2 nM Human FC-tagged human ACE2 (GenScript, Z03484). AlphaLISA detection was performed using Anti-Human IgG Alpha Donor beads (PerkinElmer, AS114) and Anti-6xHis AlphaLISA Acceptor beads (PerkinElmer, AL178). CFPS reaction containing the expressed sdFab of interest was first mixed with S trimer and allowed to incubate for 1 hour. Subsequently, ACE2 was added and allowed to equilibrate for a further 1 hour before the simultaneous addition of the acceptor and donor beads.

**Data analysis**

Because of the nature of the intended use of the reported assay as an initial screen, we use the Benjamini and Hochberg False Discovery Rate procedure[50] to correct for multiple testing, which is less conservative than other standard methods. Statistical analyses were performed in python. Two-sided t-tests were performed using the scipy package and the FDR procedure was performed using the statsmodels package with a family-wise error rate of 5%. For t-tests, the following samples were considered to be background, and the combined data were used in the t-test. Assembly: No DNA and Buffer controls. S trimer binding: No DNA and Buffer controls. RBD binding: No DNA and Buffer controls. ACE2 competition: No DNA and αHER2.
